# Stage-specific muscle wasting mechanisms in a novel cancer cachexia mouse model for ovarian granulosa cell tumor

**DOI:** 10.1101/2020.06.16.154385

**Authors:** Yaqi Zhang, Jie Zhu, So-Youn Kim, Megan M Romero, Kelly A Even, Takeshi Kurita, Teresa K Woodruff

## Abstract

Cachexia is a progressive muscle wasting syndrome that increases mortality risk in cancer patients, while there are still no effective treatment due to the complexity of syndrome and the lack of preclinical models. We identified a transgenic mice model with ovarian granulosa cell tumors mimic the progression of cachexia seen in humans, including drastic weight loss, skeletal muscle wasting and increased serum cachexia biomarker activin A and GDF15. Hypercatabolism was detected in skeletal muscle, having upregulation of E3 ligases *Atrogin-1* and *Murf-1*. Our cachexia model exhibited stage-specific muscle wasting mechanisms. At precachexia stage, elevation of activin A activates p38 MAPK. Inhibition of activin A with Follistatin reversed weight loss at precachexia stage. At cachexia stage, energy stress in skeletal muscle activates AMPKα and leads to upregulation of *FoxO3*. Our results indicate this novel preclinical cancer cachexia model is exploitable for studying pathophysiological mechanisms and testing therapeutic agents of cachexia.

## Introduction

Cachexia is a life-threatening weight loss syndrome in advanced cancer patients that severely impairs physical function, increases the severity of treatment-related toxicity, and accounts for nearly 25% of cancer-related deaths(*1*). Cancer cachexia is characterized by progressive weight loss over time, most commonly ≥5% weight loss in 6 months(*2*). Cancer cachexia is a continuum: preceding substantial body weight loss, patients with early clinical and metabolic signs (involuntary weight loss < 5%) are considered precachexia(*2*). Evidence suggests that earlier treatment of cachexia is critical, as weight loss >5% is associated with higher mortality risk(*3*), and reversing cancer cachexia is also more challenging once patients have reached advanced stages.

Most knowledge regarding the mechanisms of cancer cachexia and possible therapeutic approaches have come from preclinical animal models, including syngeneic cancer graft models, xenograft tumor models, genetically engineered spontaneous tumor models, and carcinogen-induced cancer models(*4*). Syngeneic cancer graft models, such as the C26 colon carcinoma and Lewis lung carcinoma (LLC) models, are well-established and widely used for investigation of cachexia mechanisms and drug testing. However, a major drawback of these models is the relatively short period between the onset of cachexia symptoms and death due to aggressive growth of non-spontaneous cancers. Human tumor xenograft models can closely mimic human tumor characteristics; however, the application of xenograft models to the study of cachexia is limited, primarily because immune-deficient recipient animals lack an inflammatory response, which is critical for development of cancer cachexia(*4*). Establishment of good preclinical models that develop tumors spontaneously and recapitulate tumor-host interactions are needed in order to better understand the mechanisms of cancer cachexia development and for use in drug testing.

Muscle wasting is a hallmark of cancer cachexia and is associated with increased morbidity and mortality; prevention of muscle wasting has been shown to increase survival in tumor-bearing animal models(*5, 6*). Skeletal muscle contains the largest pool of proteins in the entire organism, and is highly sensitive to energy stress under abnormal conditions. Previous studies have described muscle wasting in cancer cachexia as a loss of homeostasis between anabolic and catabolic processes in skeletal muscle(*3*), with the balance tipped toward a catabolic state as a result of activated proteolytic systems. Two muscle-specific E3 ligases, muscle-specific RING finger 1 (Murf1, also known as TRIM63) and muscle atrophy F-box (Atrogin-1/MAFbx), as well as the autophagy-related lysosomal proteolytic marker microtubule-associated protein 1 light chain 3 alpha (Lc3a), are widely used as markers of accelerated proteolysis and muscle atrophy(*7, 8*). The transcription factor Forkhead Box O3 (FOXO3) is also a key regulator in protein breakdown in muscle(*9*). FOXO3 modulates the expression of factors in the ubiquitin-proteasome and autophagy-lysosomal proteolytic pathways, including Murf-1, Atrogin-1, and Lc3a(*10–12*).

Although the development of cachexia is accompanied by an increase in inflammatory factors derived from host or tumor cells, such as interleukin (IL)-6(*13*) and tumor necrosis factor-alpha (TNF-α)(*14*), inhibiting the inflammatory response directly fails to reverse cachexia. TGF-β superfamily members such as activin A(*15, 16*), myostatin(*17*), and GDF15(*18*) have been shown to affect cancer cachexia development by negatively regulating the balance between the synthesis and degradation of muscle proteins. The soluble activin type II receptor, which blocks activin and myostatin signaling, has been shown to completely reverse muscle wasting and weight loss in cachexia mouse models(*5*). Antibodies and small molecules that target activin have also been developed(*19, 20*). In a phase 2 clinical trial, bimagrumab, a human monoclonal antibody that blocks activin type II receptors, has been shown to increase skeletal muscle mass(*21*).

As part of our research in ovarian follicle activation, we created a transgenic mouse that expresses a constitutively active allele of phosphatidylinositol-4,5-bisphosphate 3-kinase (PI3K) exclusively in oocytes(*22*). Surprisingly, these mice showed highly-penetrant (100%) transformation of the granulosa cell lineage to high-grade cancer by postnatal day (PD) 65 and succumbed to cancer cachexia by PD80(*22, 23*). This mouse model not only developed significant cancer cachexia symptoms, but also had high serum levels of activin A and GDF15, consistent with the serum profile seen in cancer cachexia patients. Here, we investigated muscle wasting progression and stage-specific mechanisms driving cachexia in our novel mouse model. We found a dominant role of activin A signaling in cachexia initiation through activation of the p38 MAPK pathway, and activation of AMPK pathway and significant upregulation of the *FoxO3* transcription factor at later stages of cachexia due to energy depletion in skeletal muscle. We concluded that our novel preclinical cancer cachexia mouse model closely mimics human cancer cachexia progression, and will be an important tool for further mechanistic studies as well as screening for molecular markers and novel therapeutics to reverse muscle wasting and metabolic disorders in patients with cancer.

## Results

### Mice with oocyte-specific constitutive PI3K activation develop systemic cancer cachexia-like symptoms and gene expression patterns in muscle wasting

Cre+ female mice with oocyte-specific constitutive PI3K activation (**Supplementary Fig. 1a**) developed bilateral granulosa cell tumors (**Supplementary Fig. 1b, c, d**) from PD60, as previously described(*22, 23*). Along with tumor formation and growth, the mice showed significant weight loss, measured starting on PD60 until 15% of their peak body weight was lost (**Fig. 1a**). All Cre+ mice (n=18) had experienced dramatic weight loss (more than 15%, defined as cachectic) and were euthanized when they reached up to 20% weight loss (**Supplementary Fig. 1e**). Weight loss started at slightly different ages: 17% (3/18) of mice showed weight loss starting around PD65 (cohort 1), 39% (7/18) around PD70 (cohort 2), 33% (6/18) around PD80 (cohort 3) and 6% (1/18) around PD85 (cohort 4) (**Fig. 1A**). Together, 100% (18/18) of Cre+ mice lost 15% of their body weight before PD100, with weight loss occurring over a period around 10 days (**Fig. 1A**). Atrophy of skeletal muscle tissue in cachectic Cre+ mice was observed by H&E staining of tibialis anterior (TA) muscles (**Fig. 1B**) and by visual examination at the time of dissection (**Supplementary Fig. 1F**). Immunostaining of the TA muscles with anti-Laminin revealed significantly smaller skeletal myofibers in cachectic Cre+ mice compared to Cre-control mice (**Fig. 1B**). Measurement of myofiber cross-sectional area (CSA) quantitatively confirmed significant atrophy of muscle fibers in Cre+ mice (**Fig. 1C**). Depletion of abdomen adipose tissue in cachectic Cre+ mice was also observed at the time of dissection (**Supplementary Fig. 1G**) and by whole-body MRI scanning (**Supplementary Fig. 1H**). Lean mass and fat mass in cachectic Cre+ mice compared with age-matched Cre-control mice were quantitatively measured by EchoMRI (**Fig. 1D**). Cachectic Cre+ mice showed additional abnormalities in various organs, including obvious cell apoptosis around the central vein in the liver, depletion of the stomach inner epithelial layer, and disarrangement of gastric glands (**Fig. 1E**). These findings are consistent with liver and stomach inner epithelial layer cell damage reported in another cancer cachexia model, inhibin-α knockout mice(*15, 24*).

**Figure 1.**
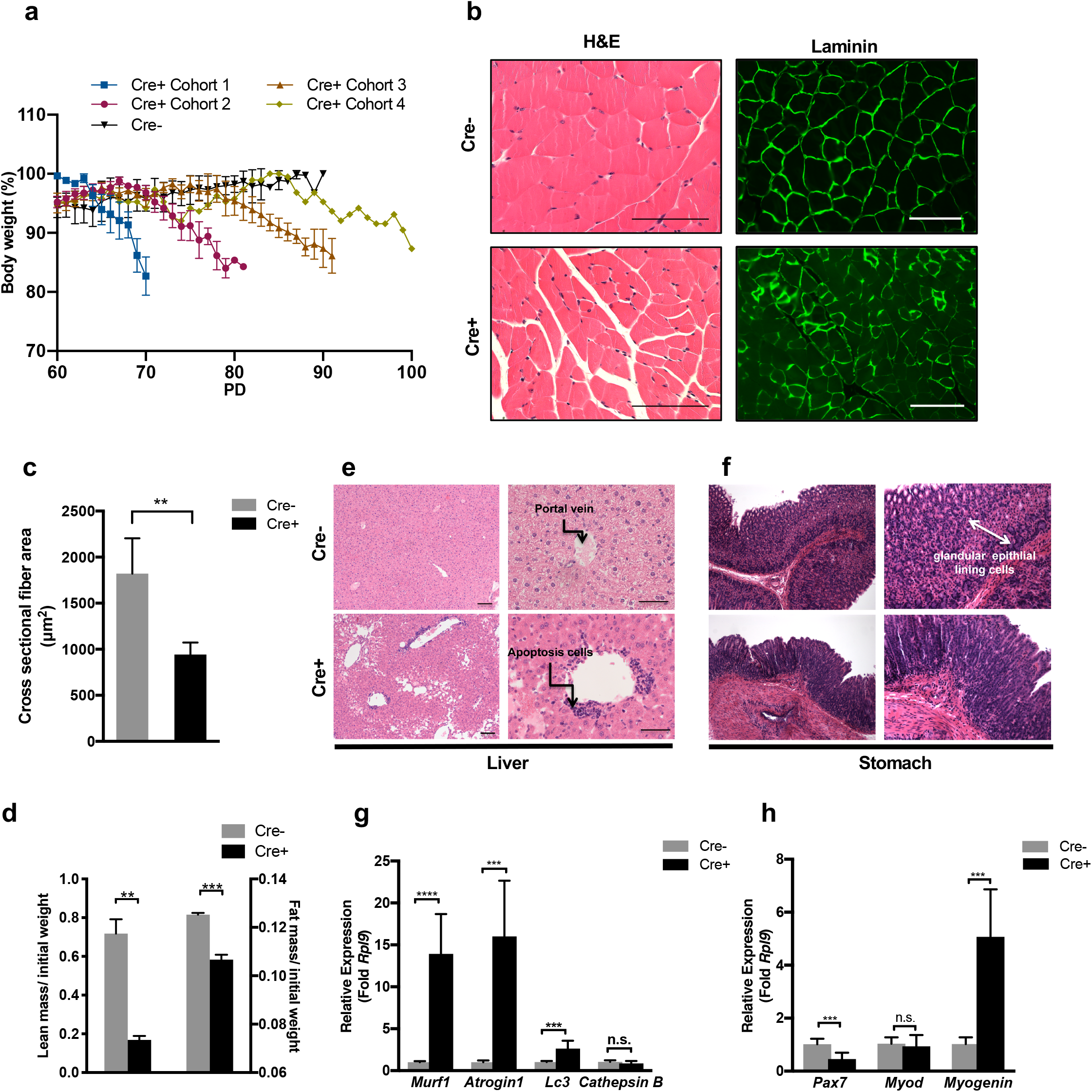
Mice with oocyte-specific constitutive PI3K activation develop systemic cancer cachexia-like symptoms. (**A**) Weight loss of cachectic Cre+ mice (n=18) and age-matched Cre-mice (n=4) from PD60. Cre+ mice were divided into 4 cohorts based on date of weight loss initiation: PD65 (cohort 1), PD70 (cohort 2), PD80 (cohort3), and PD85 (cohort 4). (**B**) Representative images of H&E staining (left) and immunofluorescence staining against Laminin (right) of skeletal myofiber bundles in cross section of TA muscle from Cre+ mouse (upper) compared with Cre-control mouse (lower). Scale bar = 20 μm. (**C**) Cross-sectional area (CSA) of myotube in TA muscle stained with Laminin from Cre+ mice and Cre-mice (n=3). (**D**) Ratio of lean mass and fat mass to initial body weight in Cre+ mice with 15% weight loss (n=3) and age-matched Cre-mice (n=3) using EchoMRI. (**E**) Representative images of liver histology for age-matched Cre-mice and Cre+ mice with 15% weight loss. The portal vein (black arrow) in Cre+ mice is surrounded by a large number of apoptotic cells compared with Cre-mice. Scale bar = 100 μm or 20 μm. (**F**) Representative images of stomach histology for Cre+ mice with 15% weight loss and age-matched Cre-mice. The glandular epithelial lining cell layer in the Cre-mouse is marked by the white double arrow line. Scale bar = 100 μm or 20 μm. (**G**) RT-PCR analysis of *Murf1*, *Atrogin1*, *Lc3a*, and *Cathepsin B* in tibialis anterior (TA) muscle from Cre+ mice with 15% weight loss and age-matched Cre-mice (n=8). (**H**) RT-PCR analysis of *Pax7*, *MyoD1*, and *Myogenin* in TA muscle from Cre+ mice with 15% weight loss (n=8) and age-matched Cre-mice (n=8). Data are shown as mean ± s.d. of biological replicates. *, **, ***, and **** represent *P*<0.05, *P*<0.01, *P*<0.005, and *P*<0.001, respectively.

Given that skeletal muscle wasting is a hallmark of cachexia, we quantified the expression of genes known to be involved in this process by RT-qPCR in the TA muscle from cachectic Cre+ and age-matched Cre-control mice: muscle-specific E3 ligases *Atrogin-1* and *Murf-1*, the lysosomal proteolytic factors *Lc3a* and *Cathepsin B*, as well as the calcium-dependent proteolytic factor *Calpain 13*. Transcripts for *Atrogin-1*, *Murf-1*, and *Lc3a* were significantly enriched in the TA muscle from cachectic Cre+ mice (**Fig. 1G**), indicating strong activation of the ubiquitin-proteasome system and autophagy. Other muscle degenerative pathways were not activated; for example, *Cathepsin B* mRNA levels were not different in Cre+ and Cre-TA muscle tissue (**Fig. 1G**) and the levels of *Calpain 13* mRNA were below the limit of detection (data not shown). The caspase system was not activated in the TA muscle of Cre+ mice, as indicated by the low levels of *caspase 3* expression in the Cre+ mice (**Supplementary Fig. 2A**). Expression of genes known to be involved in skeletal muscle regeneration were also measured. The transcript for muscle satellite cell activation factor *Pax7* was significantly lower in Cre+ mice compared to Cre-mice (**Fig. 1H**), indicating the depletion of muscle satellite cells. The expression of myoblast proliferation factor *MyoD1* was not different, whereas expression of the myoblast differentiation factor *Myogenin* was significantly higher in TA muscle from cachectic Cre+ mice compared to Cre-controls (**Fig. 1H**), suggesting that muscle regeneration was not impaired. Overall, the gene expression patterns indicate that activation of the ubiquitin proteasome system and autophagy are major mechanisms that induce muscle atrophy in cachectic Cre+ mice.

### Cre+ mice have elevated serum cachectic biomarkers during cachexia progression

Skeletal muscle wasting in cancer cachexia is associated with an increase in various factors that derive from host or tumor cells. A previous study in which various cytokines were measured in patients at different stages of cachexia reported a significant increase in two TGFβ family members, activin A and GDF15, starting from early-stage (precachexia) disease, that remained high as the disease progressed(*18*). Therefore, we tested serum levels of activin A and GDF15 in our cachectic Cre+ mice. The serum level of activin A in cachectic Cre+ mice was elevated almost 100-fold compared to age-matched Cre-mice, while GDF15 was more than 10-fold higher compared to age-matched Cre-mice (**Fig. 2A**). We measured serum levels other inflammatory factors previously reported to be involved in the development of cancer cachexia, such as IL-1β, IL-6 and TNF-α(*13, 14, 25*). IL-6 and TNF-α, but not IL-1β, were significantly higher in cachectic Cre+ mice compared to age-matched Cre-mice (**Fig. 2B**).

**Figure 2.**
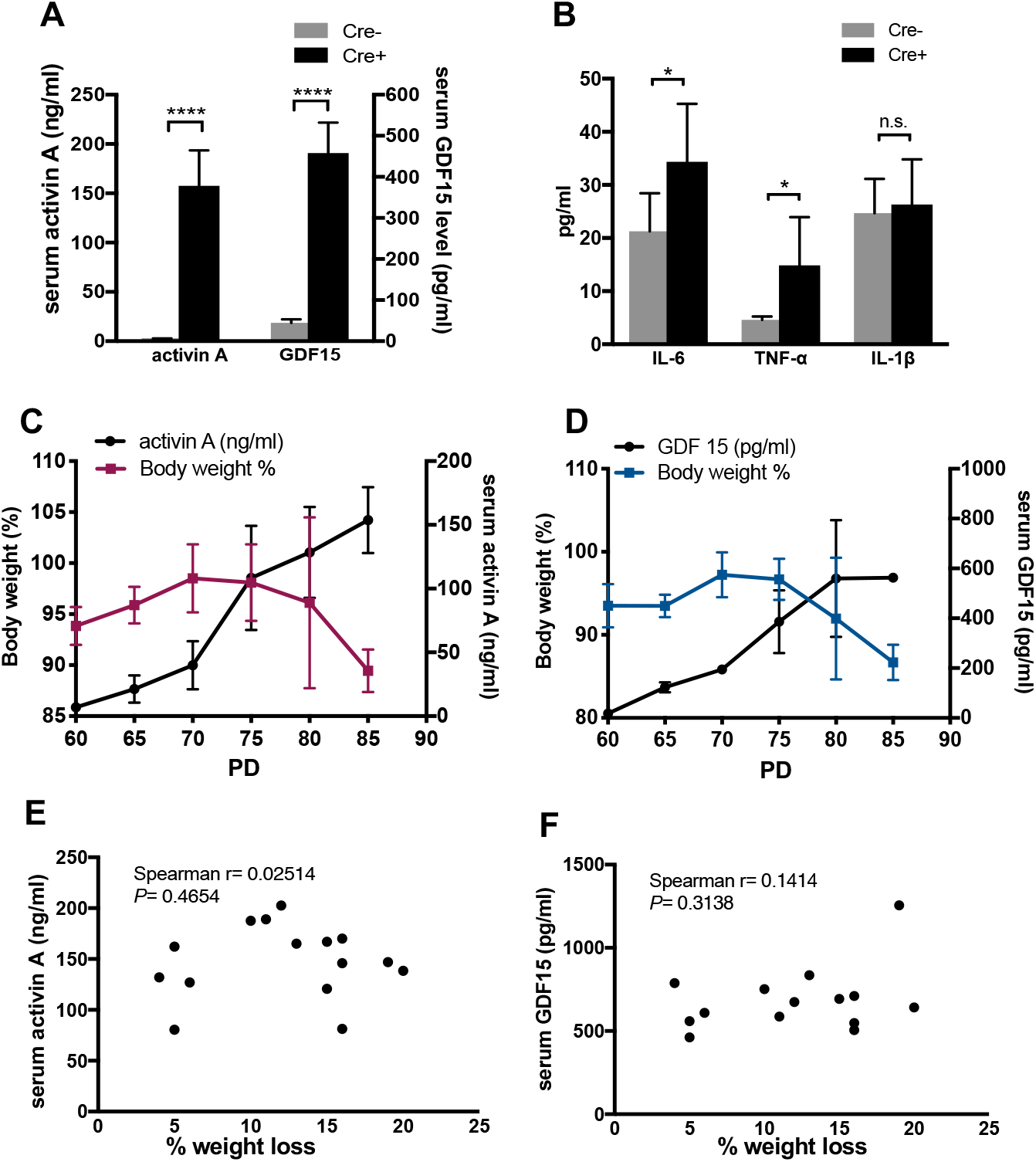
Weight loss in cachexia mice is correlated with serum activin A and GDF15 levels. (**A**) Serum activin A and GDF15 levels in Cre+ mice (n=9) with 15% weight loss based on their peak weight and age-matched Cre-mice (n=9) measured by ELISA. (**B**) IL-6, TNF-α, and IL-1β level in Cre+ mice (n=6) with 15% weight loss and age-matched Cre-mice (n=6) measured by ELISA. (**C**) Weight and activin A serum levels from tail blood draws in Cre+ mice (n=5) starting on PD60 until mice had 15% weight loss based on the peak weight at different time points. (**D**) Weight and GDF15 serum levels from tail blood draws in Cre+ mice (n=4) starting on PD60 until mice had 15% weight loss based on the peak weight at different time points. (**E**) Correlation between serum activin A levels with weight loss from 5% to 20% in Cre+ mice (n=15) and (**F**) correlation between serum GDF15 levels and weight loss from 5% to 20% in Cre+ mice (n=15). “r” represents correlation value and Spearman correlation used for the statistical analysis to calculate p value. Data are shown as mean ± s.d. of biological replicates. * and **** represent *P* <0.05 and *P*<0.001, respectively.

To establish the relative timing and degree of weight loss to the serum levels of activin A and GDF15, we measured serum activin A (n=5) and GDF15 (n=4) in tail blood draws from Cre+ mice starting on PD60 until the mice reached 15% weight loss based on the peak weight. Cre+ mice (n=15) were euthanized at different levels of weight loss from 5% up to 20% and the serum level of cytokines was determined. Serum levels of activin A (**Fig. 2C**) and GDF15 (**Fig. 2D**) were significantly elevated in Cre+ mice before evidence of body weight loss. Levels of activin A (**Fig. 2E**) and GDF15 (**Fig. 2F**) serum level did not further increase as weight loss began, suggesting their potential role in the initial stages of the disease. Serum levels of TNF-α also did not show a strong correlation with continuous weight loss (**Supplementary Fig. 2B**). Together, the circulating cytokines that were elevated in the precachexia stage remained relatively stable level during progression of cachexia.

### Activin A/p38 MAPK signaling is activated in skeletal muscle at the precachexia stage

Given that activin A negatively regulates muscle, the high activin A in Cre+ mice serum may play an important role in initiating muscle wasting. To test this hypothesis, we treated Cre+ mice with recombinant Follistatin Fst288, a native activin bio-neutralizing protein(*26*). Three Cre+ mice were treated with PBS or Fst288 at two different time points: the precachexia stage (weight loss up to 5%) and cachexia stage (continuous weight loss between 5%-10%). Treatment at the precachexia stage was stopped when body weight returned to baseline, then the disease was allowed to progress until the cachexia stage and the second treatment time point. As expected, all three Cre+ mice that received PBS injection continued losing weight until 15% weight loss (**Supplementary Fig. 3A**). Fst288 injection during the first period was 100% effective, as it reversed weight loss in all three precachectic Cre+ mice (**Fig. 3A**, **Supplementary Fig. 3B**). Fst288 injection during the second period in same set of three Cre+ mice during cachexia stage was less effective: only one Cre+ mouse that was injected with Fst288 starting at 10% weight loss showed a reversal in weight loss (**Fig. 3B**), whereas the other two Cre+ mice had minimal response to Fst288 and continued to lose weight (**Fig. 3B**). The initial reversal of weight loss with Fst288 treatment, which extended survival in Cre+ mice treated with Fst288 compared with Cre+ treated with PBS, implies that high serum levels of activin A may be involved in the initial weight loss in cachexia.

**Figure 3.**
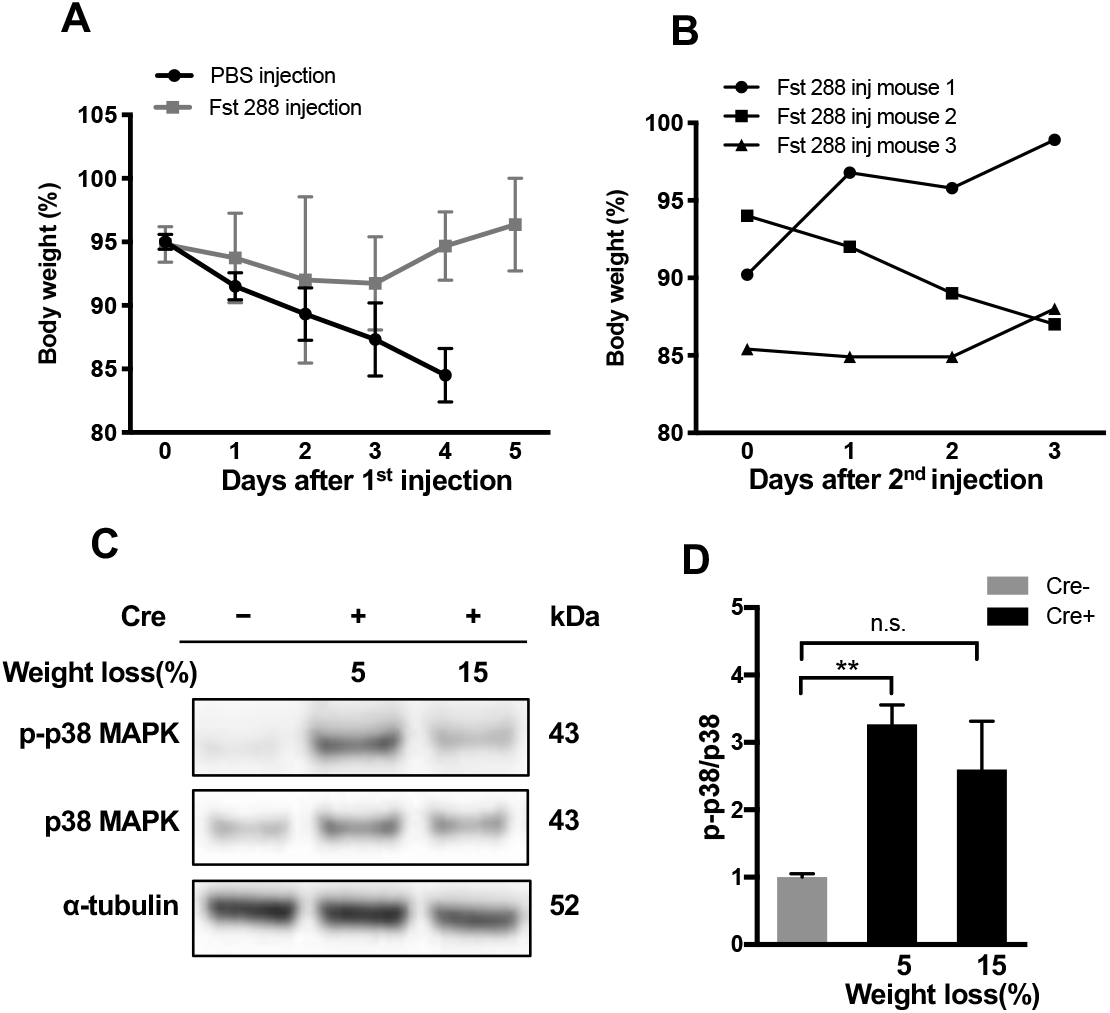
Activin A/p38 MAPK signaling is activated in skeletal muscle of mice at the precachexia stage and initiation of weight loss. (**A**) Weight loss (%) of Cre+ mice (n=3) with PBS solution injection or Cre+ mice (n=3) injected with Fst288 200 μg/kg twice/day starting at less than 5% weight loss. (**B**) Weight loss (%) of three individual Cre+ mice that received Fst288 injection during the early stage of weight loss, then Fst288 200 μg/kg twice/day from 5-15% weight loss. (**C**) Representative immunoblot of p-p38 MAPK, total p38 MARK, and α-tubulin in TA muscle protein lysates collected from Cre+ mice with 5% weight loss (n=3) and 15% weight loss (n=3) and Cre-mice (n=3) age-matched to Cre+ mice with 15% weight loss. (**D**) Quantification of densitometry analysis for immunoblot images for the expression protein ratio of p-p38/p38 proteins from TA muscle collected from Cre+ mice with 5% and 15% weight loss and age-matched Cre-mice (n=3) using ImageJ software. Data are shown as mean ± s.d. of biological replicates. Non-paired t-test was used for statistical analysis. * and ** represent *P*<0.05 and *P*<0.01, respectively.

To further examine the downstream targets of activin A in muscle, we investigated the p38β mitogen-activated protein kinase (MAPK) pathway, which has been shown to respond to activin A stimulation and induce transcriptional expression of E3 ubiquitin ligases in skeletal muscle(*27*). Phospho-p38 MAPK was highly activated in the TA muscle from precachectic Cre+ mice (≤5% weight loss) (**Fig. 3C**, **D**), but decreased to baseline in cachectic Cre+ mice (15% weight loss). There was no significant difference in phospho-p38 MAPK/p38 MAPK in cachectic Cre+ mice and age-matched Cre-control mice (**Fig. 3C, D**), suggesting that the p38 MAPK signaling does not play a major role once mice have progressed past the precachectic stage. In contrast, the expression of Murf-1, Atrogin-1, and Lc3 increased continuously with disease progression and showed significant positive correlation with weight loss (**Supplementary Fig. 3C, D, E**). This suggests that other muscle wasting mechanisms, beyond activin A induction of p38 MARK signaling, likely become dominant in later stages of cachexia. These results are consistent with our observation that weight loss in Cre+ mice could not be reversed by Fst288 treatment once the mice reached 10% weight loss.

### Upregulation of *FoxO3* expression by AMPKα is dominant at later cachexia stages

Given that activin A/p38 MAPK signaling was only active at the precachexia stage in our Cre+ mice, while the expression of Murf-1, Atrogin-1, and Lc3 increased with continuous weight loss, we posited that other cachectic factors contribute to progressive weight loss in later stages of the disease. FOXO3 is one of the most important transcription factors that modulates muscle energy homeostasis and controls muscle protein breakdown by regulating E3 ubiquitin systems and the autophagy-lysosomal proteolysis system(*9*). Therefore, we examined the expression and functions of FOXO3 in our cachexia mouse model. Expression of *FoxO3* in the TA muscle from Cre+ mice was strongly correlated with the degree of weight loss (**Fig. 4A**). As expected, the transcriptional level of *Murf1*, an E3 ligase and transcriptional target of FOXO3, was positively correlated with the expression of *FoxO3* in the TA muscle from Cre+ mice with 5% up to 20% weight loss (**Fig. 4B**). FOXO3 protein levels were significantly higher in TA muscle from cachectic Cre+ mice (15% weight loss) compared with Cre-mice (**Fig. 4C, D**). Notably, FOXO3 protein was not elevated in the TA muscle from precachectic Cre+ mice (≤5% weight loss), indicating that upregulation of FOXO3 is more dominant at later stages of the disease (**Fig. 4C, D**). Phosphorylation of FOXO3 on Thr32 has high in TA muscle tissue from both precachectic and cachectic Cre+ mice compared with age-matched Cre-mice (**Fig. 4C, D**). The ratio of phospho-FOXO3/FOXO3, which reflects the ratio of inactivated FOXO3 in cytoplasm to total cellular FOXO3 protein, was higher in TA muscle from both precachectic and cachectic Cre+ mice compared with age-matched Cre-mice, with no significant difference between the precachexia and cachexia mice (**Fig. 4E**). To detect the nuclear translocation of FOXO3, TA muscle from precachectic and cachectic Cre+ mice and age-matched Cre-mice were stained with FOXO3 antibody (**Fig. 4F**). Nuclear staining of FOXO3 and overall cellular levels of FOXO were higher in TA muscle of cachectic Cre+ mice, whereas nuclear staining of FOXO3 was lower in TA muscle of precachectic Cre+ mice. Cytoplasmic FOXO3 was only slightly higher in TA muscle of precachectic Cre+ mice compared with Cre-mice. These findings indicate that an increase in FOXO3 nuclear translocation occurs in later stages of cachexia.

**Figure 4.**
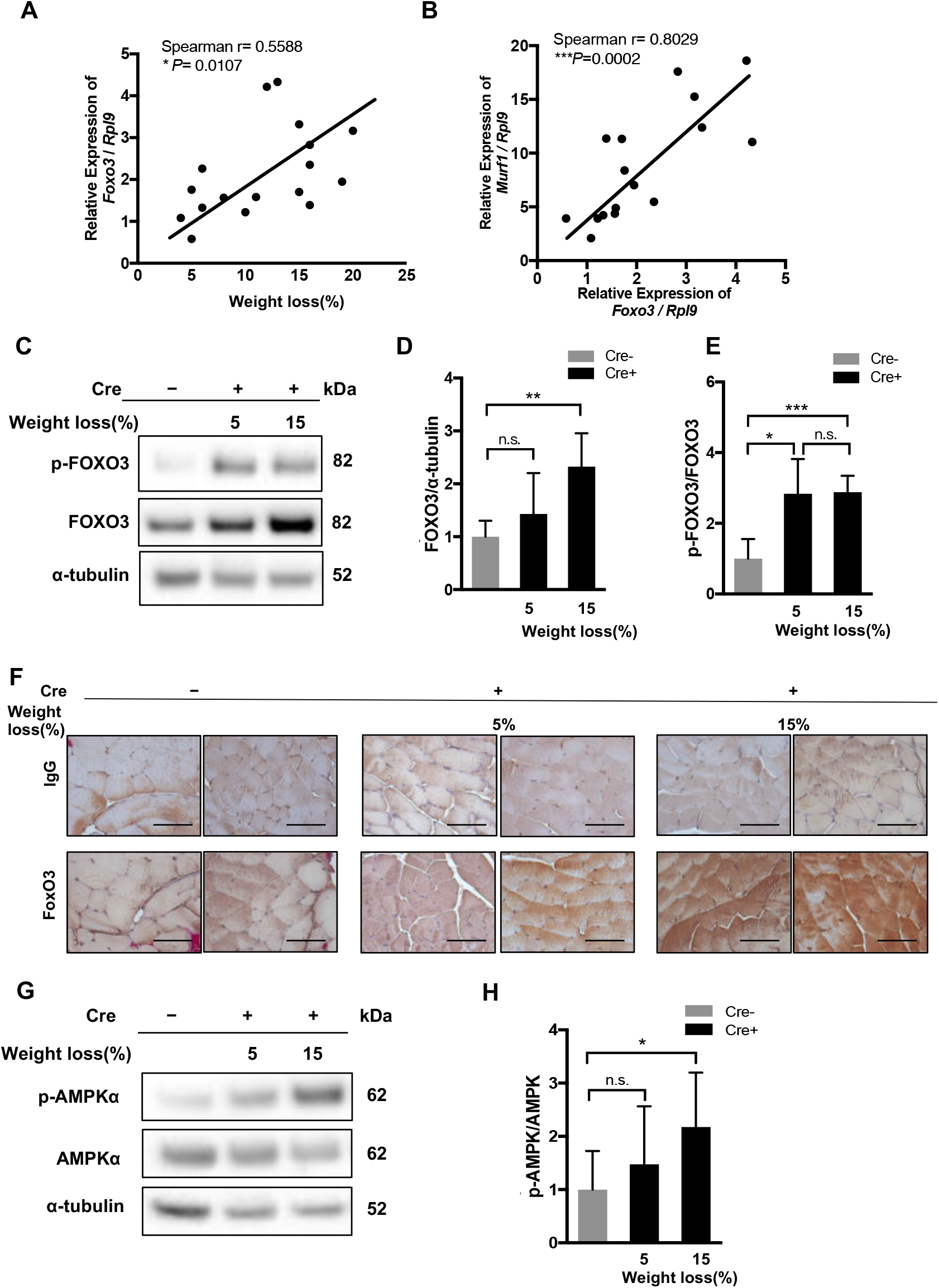
*FoxO3* upregulation by AMPKα signaling mediates muscle wasting in late stages of cachexia. (**A**) Correlation between *FoxO3* mRNA expression in TA muscle and percentage of weight loss in Cre+ mice (n=17) and (**B**) correlation between *Murf1* and *FoxO3* mRNA expression in TA muscle from Cre+ (n=16) graphed using Spearman correlation by Prism 7 software. (**C**) Representative immunoblots of p-AKT, AKT, p-FOXO3, and FOXO3 protein from TA muscle lysate collected from Cre+ mice with 5% weight loss, Cre+ mice with 15% weight loss, and age-matched Cre-mice. (**D-E**) Quantification of densitometry analysis for immunoblot images for the expression protein ratio of (**D**) total FOXO3/ α-tubulin, (**E**) p-FOXO3/FOXO3 in Cre+ mice with different weight loss (n=3) and age-matched Cre-mice (n=3) using ImageJ software. (**F**) Representative images of immunohistochemistry staining against FOXO3 in TA muscle from Cre+ mice (n=2). IgG was used as negative control Scale bar = 20 μm. (**G**) Immunoblot analysis of AMPKα, p-AMPKα, and α-tubulin protein for TA muscle lysate collected from Cre+ mice with 5% weight loss, Cre+ mice with 15% weight loss, and age-matched Cre-mice. (**H**) Quantification of densitometry analysis of immunoblot images for ratio of p-AMPKα/ AMPKα in Cre+ mice with 5% weight loss, Cre+ mice with 15% weight loss, and age-matched Cre-mice (n=3). Non-paired t-test was used for statistical analysis by Prism 7 software. Data are shown as mean ± s.d. of biological replicates. * and ** represent *P*<0.05 and *P*<0.01, respectively.

Adenosine monophosphate-activated protein kinase (AMPK) has been found to stimulate muscle protein degradation by increasing FOXO3 transcription, thereby stimulating the expression of *Atrogin-1* and *Murf-1*(*28*). As AMPK is a highly conserved master regulator of metabolism and a sensor of energy stress(*29*), we examined AMPK levels and activation in TA muscle from Cre+ mice at different stages of weight loss. Phospho-AMPK was not different in TA muscle from precachectic Cre+ mice compared with age-matched Cre-mice, but was dramatically higher in TA muscle from cachectic Cre+ mice (**Fig. 4G**). Consequently, the ratio of phospho-AMPK/AMPK was not significantly different between precachectic Cre+ and age-matched Cre-mice, while the ratio was significantly higher in cachectic Cre+ mice (**Fig. 4H)**. The activation of AMPK in cachectic Cre+ mice implies an increase in energy stress in TA muscle as mice progress to later stages of the disease.

To determine whether energy stress upregulates FOXO3 expression or induces its activation, we measured the expression of *FoxO3* in TA muscle tissue from two other mouse models of energy stress, a fasting model and type I diabetes model (**Supplementary Fig. 4**). After a 16-hour fast, expression of *FoxO3* in TA muscle was significantly elevated (**Supplementary Fig. 4A**) compared with that of non-fasting mice. Expression of *FoxO3* in muscle from mice with type 1 diabetes was also higher than in non-diabetic controls (**Supplementary Fig. 4B**). Total FOXO3 protein levels were significantly higher in both fasting mice and diabetic mice compared to fed and non-diabetic controls, respectively (**Supplementary Fig. 4C, E, G**); this was comparable to the pattern seen in cachectic Cre+ mice. The phospho-FOXO3/FOXO3 ratio was not significantly different in normal fed mice and fasting mice, due to a simultaneous increase in both phospho-FOXO3 and total FOXO3 protein (**Supplementary Fig. 4D**), again similar to the changes seen in cachectic Cre+ mice. The significantly higher total FOXO3 protein level in the TA muscle tissue of the type 1 diabetes model led to a much lower ratio of phospho-FOXO3/FOXO3 compared with control mice (**Supplementary Fig. 4F**). The similarity in molecular phenotypes between cachectic Cre+ mice and the fasted and type 1 diabetes models support the notion that Cre+ mice develop energy stress in muscle in the later stages of disease progression.

### Hypermetabolism and tumor-induced energy depletion promote the development of cachexia

Cancer cachexia is a complex metabolic disorder marked by dramatic changes in energy balance. We hypothesized that energy stress and energy depletion are the major factors that cause muscle wasting at later stages of cachexia in Cre+ mice. To evaluate the energy available to muscle tissue, we measured glucose metabolism status in our cachexia mouse model. Although a previous study found evidence of insulin resistance in cancer cachexia(*30*), we found higher levels of insulin receptor and activation of phospho-AKT in TA muscle from Cre+ mice compared to Cre-controls (**Supplementary Fig. 5A**), indicating that the muscle is still sensitive to glucose and that energy stress in the Cre+ mice was not due to insulin resistance. Thus, we next examined non-fasting blood glucose level in precachectic Cre+ mice (≤5% weight loss) and cachectic Cre+ mice (15% weight loss) compared with age-matched Cre-mice. Both precachectic and cachectic Cre+ mice similarly had significantly lower non-fasting blood glucose levels (around 100 mg/dL) compared to Cre-mice (around 170 mg/dL) and previously reported normal C57BL/6 mice(*31*) (**Fig. 5A**). In contrast, fasting blood glucose in the cachectic Cre+ mice (15% weight loss) was dramatically lower (around 35 mg/dL) than age-matched Cre-mice and precachectic Cre+ mice, indicating hypoglycemia in cachectic Cre+ mice; while precachectic Cre+ mice maintained a fasting blood glucose level around 70-75 mg/dL, although it was still significantly lower than that in the Cre-mice (90 mg/dL) (**Fig. 5B**). We also performed a glucose challenge in the Cre+ and age-matched Cre-mice; oral fed glucose was cleared within 2 hours in all of the groups (**Fig. 5C**), with a peak at 15 min after feeding and a gradual decrease to the normal range by approximately 90 minutes. However, the peak glucose value in the cachectic Cre+ mice was lower than that seen in the precachectic Cre+ mice or the Cre-mice (**Fig. 5C**). Overall, these studies suggest that blood glucose metabolism is significantly altered in later stages of cachexia in our Cre+ mouse model. To map the distribution of glucose against progressive weight loss in the Cre+ mice, we monitored glucose usage in different tissues with ^18^F-fluro-deoxyglucose (^18^F-FDG) positron emission tomography-computed tomography (PET-CT). Regional uptake of ^18^F-FDG reflects the activity of various glucose transporters and hexokinase in manner similar to glucose, allowing us to characterize differences in glucose consumption. ^18^F-FDG PET/CT was used to measure glucose consumption in Cre+ mice at three different disease stages: tumor formation stage (PD55) (n=3), precachexia stage (PD65) (n=3), and cachexia stage (15% weight loss) (n=1). Age-matched Cre-mice were used as the control group (**Fig. 5D**). Only one Cre + mouse had been monitored longitudinally from tumor initiation stage all the way to cachectic stage due to the weak body condition of cachectic mice. At the tumor initiation stage, no ^18^F-FDG uptake was observed in the Cre+ mouse abdomen, although tumor could be detected by MRI (**Fig. 5D**). At the precachexia stage, one clear area of ^18^F-FDG uptake was seen in the tumor region; in the cachectic Cre+ mouse, multiple areas of ^18^F-FDG uptake were observed in the tumor region, indicating elevated glucose consumption (**Fig. 5D**). This hypermetabolism within the tumor tissue may contribute to the energy stress in muscle tissue, leading to wasting. In addition to elevated glucose consumption in tumor tissue, the brown adipose tissue, located in the neck area of mice, also showed elevated ^18^F-FDG uptake in both Cre- and Cre+ mice (**Fig. 5D**). The signal in brown adipose tissue in precachectic Cre+ mice (n=3) was slightly higher compared with age-matched Cre-mice (**Supplementary Fig. 5B**), but the signal varied widely across the different Cre+ mice. ^18^F-FDG uptake in brown fat was dramatically lower in the cachectic Cre+ mouse, indicating depletion in all adipose tissue (**Fig. 5D**). Taken together, the distribution of ^18^F-FDG suggests that the glucose consumption of tumor tissue increases as tumor grows, contributing to the energy stress of cachectic Cre+ mice muscle, which has less glucose available.

**Figure 5.**
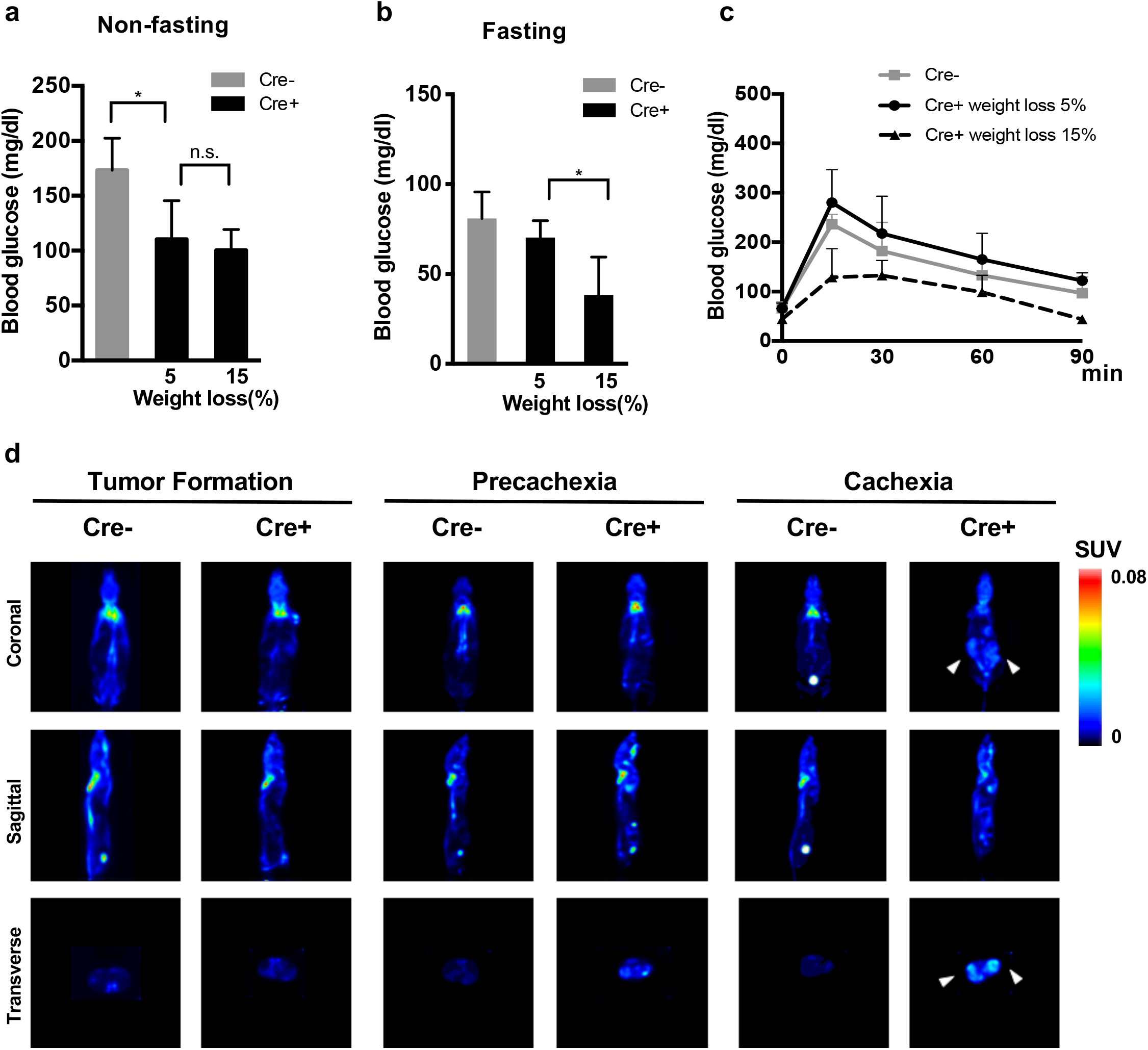
Hypermetabolism and tumor-induced energy depletion in skeletal muscle in the cachexia mouse model. (**A**) Glucose level in normally fed Cre+ mice with 5% weight loss and 15% weight loss compared with age-matched Cre-mice (n=5). (**B**) Glucose level after fasting for 16h in Cre+ mice with 5% weight loss and 15% weight loss compared with Cre-mice (n=4). (**C**) Glucose challenge in Cre+ mice with 5% weight loss, Cre+ mice with 15% weight loss, and Cre-mice age-matched with Cre+ mice with 15% weight loss (n=3). The blood glucose level was measured before 2 mg/kg glucose feeding and 15, 30, 60, and 90 mins after glucose feeding. (**D**) Representative images of distribution of ^18^F-FDG in brown adipose tissue (red arrows) and tumor (white arrows) in same Cre+ mouse from three different stages: tumor initiation stage, weight stable stage, and over 15% weight loss stage compared with age-matched Cre-mouse using PET/CT scan. The representative images were normalized by the same scan location and density. Non-paired t-test was used for statistical analysis by Prism 7 software. Data are shown as mean ± s.d. of biological replicates. * and ** represent *P*<0.05 and *P*<0.01, respectively.

## Discussion

Cancer cachexia is a multifactorial process caused by complex pathogenic factors that involve both tumor and host tissue. In this study, we characterized an ovarian granulosa cell tumor (GCT) mouse model with spontaneously developed cancer cachexia symptoms. We found that this cachexia mouse model not only develops weight loss and muscle wasting as other cachectic models, but also has a similar plasma cytokine profile as cancer cachexia patients. Most importantly, we demonstrated that this GCT cachexia model has stage-specific signaling associated with muscle wasting, with activin A/p38 MAPK signaling dominant at the precachexia stage that switches to energy stress/AMPK/FOXO3 signaling at a later cachectic stage. We examined the involvement of plasma cytokines and energy stress in cachexia development in our novel mouse model.

With complete GCT penetrance in as previous reported(*23*), all Cre+ mice exhibited cachexia symptoms, including continuous weight loss, abnormal pathology in the liver and stomach, loss of adipose tissue, and most importantly, significant wasting in muscle tissue (**Fig. 1**). Although the onset of cachexia-associated weight loss varied with time, occurring primarily from PD65 to PD80, we found an association between an increase in serum levels of activin A and GDF15 and onset of weight loss. The cytokine profile of Cre+ mice showed continuous elevation of activin A and GDF15 prior to the initiation of weight loss (precachexia stage, weight loss ≤5%) and consistently elevated activin A and GDF15 maintained during progressive weight loss (cachexia stage, weight loss 15%) (**Fig. 2**); this pattern is consistent with reported cytokine screening results from cancer cachexia patients(*18, 32*). The high level of serum activin A results from supraphysiological levels of activin A produced by the fast-growing granulosa cell tumors, as revealed in the previous study on this mouse model, which stimulate tumor growth in an autocrine manner(*23*). GDF15 signaling has been linked to anorexia by interacting with GDNF family receptor α-like (GFRAL) in the brain(*33*); however, the mechanisms of GDF15 in cachexia development in our Cre+ mice remain unclear.

Muscle wasting is one of the major features of cancer cachexia syndrome that seriously impairs patient quality of life. In our cachexia model, we also observed significant muscle wasting (**Fig. 1D, Supplementary Fig. 1F**). We examined possible catabolic and anabolic wasting mechanisms in the skeletal muscle, including the lysosomal proteolytic system, calcium-dependent proteolytic system, cysteine-dependent aspartate specific proteases, and the E3 ubiquitin-ligase system. In our cachexia model, we detected increased expression of molecules associated with the E3 ubiquitin-ligase system, such as *Murf1* and *Atrogin1*, and the lysosomal proteolytic system, such as *Lc3a* (**Fig. 1G, F**). Similar expression patterns have been reported in other cancer cachexia mouse models, such as inhibin-α knockout mice and syngeneic cancer graft models with C26 colon carcinoma, Lewis lung carcinoma (LLC), and Yoshida AH-130 hepatoma(*5, 34*). Similar to patients with cachexia, *caspase 3* was not upregulated/activated in our model system (**Supplementary Fig. 5B**)(*35*). We also found that p-AKT was significantly elevated in the TA muscle of cachectic Cre+ mice, suggesting that muscle atrophy in cancer cachexia is not due to the blockade of protein synthesis (**Supplementary Fig. 5**). Transcription of *Myogenin*, which is activated by p-AKT in skeletal muscle (*36*), was also elevated. Expression of myoblast proliferation factor *MyoD1* was unchanged in cachectic Cre+ mice. Our data suggest that our cachexia mouse model has relatively normal muscle regeneration activity, including myoblast proliferation and differentiation. The activation of muscle protein synthesis pathways in the cachectic mice could be a compensatory response to the acute loss of muscle fibers. These results agree with previous findings that inhibition of protein synthesis was not sufficient to induce muscle wasting in mice and humans(*37*). However, we cannot completely exclude a contribution of defects in muscle satellite cell activation to muscle wasting in our cancer cachexia model, as satellite cell marker *Pax7* was downregulated, which may signal depletion of the muscle satellite cell reserve.

Activin A has been reported as an important driving factor of cachexia in multiple previous studies, which showed that weight loss could be reversed through blocking activin signaling(*5, 16*). Cachexia symptoms are also seen in an inhibin-α knockout mouse, which also expresses supraphysiological levels of activin A(*38*). In our cachexia mouse model, supraphysiological levels of activin A appear to drive weight loss at early stages of cachexia. Serum activin A level increased along with weight loss initiation and remained high during cachexia development (**Fig. 2C**). Furthermore, neutralizing activin A by recombinant Follistatin, Fst288 partially reversed weight loss in Cre+ mice and prolonged their survival (**Supplementary Fig. 3B**). Activin A was previously found to stimulate the transcription of E3 ligase *Murf-1* and lysosomal proteolytic factor *Lc3* through activation of p38β MAPK in skeletal muscle(*27*). We confirmed that the activin A/p38 MAPK pathway is strongly activated in the TA muscle in our Cre+ mice at the precachexia stage (<5% weight loss), supporting the model that high levels of circulating activin A cause of strong catabolism in skeletal muscle. However, we also observed that p38 MAPK activation decreased as weight loss continued in cachexia progression, suggesting that factors other than activin A are likely involved in muscle wasting. This is consistent with the findings of our experiment in which Fst288 treatment reversed weight loss in the precachectic stage but was less effective in the later cachectic stage (**Fig 3A, Supplementary Fig. 3B**), indicating that blocking activin A alone is not sufficient to impede weight loss as the disease progresses. Other factors, such as energy stress and GDF15, became more dominant factors in these later stages. Thus, therapeutic use of activin A antagonists may be most beneficial at precachexia stage. Further experiments are needed to test whether higher concentrations or more frequent dosing of Fst288 or other activin A antagonists can prevent cachexia-related weight loss.

Metabolic dysfunction has been increasingly emphasized as a contributing factor in cachexia in recent years. Previous work suggests that FOXO3 transcription factors can be activated in skeletal muscle under conditions of energy stress such as starvation and type 1 diabetes(*11, 39*). Consistent with this work we found that *FoxO3* mRNA levels were elevated in both of these energy stress models and that *FoxO3* expression and FOXO3 activation was increased in our mouse model at later stages of cachexia, with a strong positive correlation with the degree of weight loss. In contrast, FOXO3 activation at the precachexia stage was not different from that in Cre-control mice, indicating that FOXO3 is not a direct target of activin A/p38 MAPK signaling. The activation of FOXO3 at later stage of cachexia suggests that FOXO3 becomes the dominant factor involved in muscle wasting as a result of energy stress, and that this occurs through direct upregulation of *FoxO3* mRNA. This mechanism is different from many cancer cachexia mice models such as C26 tumor bearing mice, where FOXO3 regulates the ubiquitin system and autophagy system through post-translational modification such as phosphorylation inhibition(*5*). AMPK, a sensor of cellular energy depletion, may stimulate *FoxO3* expression and induce skeletal muscle protein degradation through Murf1 and Atrogin-1(*28*). We demonstrated that AMPK is strongly activated in the later stages of cachexia in our model (**Fig. 4G**), providing further support that energy stress-induced *FoxO3* upregulation is the dominant pathway that promotes muscle wasting in our cachexia model.

Energy stress, especially increased resting energy expenditure due to the tumor burden and/or white fat “browning,” has been strongly associated with weight loss in cachexia patients(*40*). We characterized glucose metabolism in Cre+ mice by measuring blood glucose levels and using ^18^FDG-PET/CT scanning to track the distribution of glucose consumption. We found that the fast-growing tumor tissue showed increased glucose uptake (**Fig. 5D, lower panel**), which may account for energy depletion and energy stress in skeletal muscle. We also observed that brown adipose tissue had trend of higher glucose uptake in the precachexia stage than in the later cachexia stage (**Fig. 5D, upper panel**), suggesting overactivated brown adipose tissue as another source of glucose consumption. Further studies are needed to understand whether white fat “browning” is a contributing factor to the progression of energy stress in cachexia.

We also found that fasting blood glucose levels in Cre+ mice at the precachexia stage were comparable to that of Cre-mice (**Fig. 5B**), suggesting that precachectic mice can accommodate the increased energy consumption through gluconeogenesis, thereby maintaining energy homeostasis in the skeletal muscle. As tumor growth and hypermetabolism increases tumor glucose consumption, and the loss of adipose tissue further impairs gluconeogenesis, fasting glucose levels significantly decreased in mice at later stages of cachexia. Consequently, the skeletal muscle is deprived of glucose, which exacerbates energy stress in skeletal muscle and leads to wasting and weight loss as cachexia progresses to later stages.

Finally, although glucose resistance has been reported in cachexia patients(*30*), the glucose challenge experiment in showed that Cre+ mice have rapid glucose clearance even at the cachexia stage. This implies that insulin resistance is not the cause for energy stress in the muscle tissue in our mice model.

In conclusion, a preclinical model that mimics the complexity of human cancer cachexia development could be an effective tool for studying the mechanisms in muscle wasting, identifying novel biomarkers and therapeutic targets, and testing potential therapies. Here, we characterized a novel transgenic spontaneous developed cancer cachexia mouse model in which an activin A/p38 MAPK pathway is associated with initiation of cachexia symptoms, and the progression of weight loss is associated with *FoxO3* upregulation and energy stress due to hypermetabolism of tumor tissue and possible adipose tissue browning (**Fig. 6**). These mice provide a valuable model to track and study the development of cancer cachexia in different stages, which can be studied to better understand the pathophysiological mechanisms of cachexia and for testing therapeutics that targeting key factors such as activin A and GDF15. Furthermore, our study indicates the importance of early intervention at the precachexia stage to prevent energy stress in skeletal muscle.

**Figure 6.**
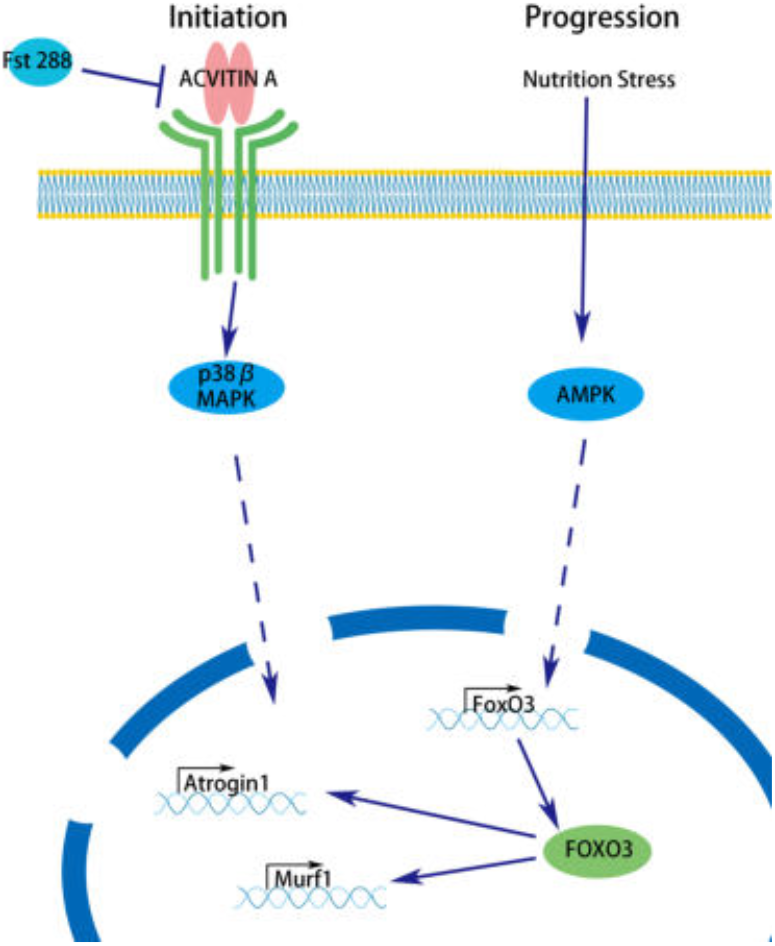
Schematic illustration of the mechanisms driving cancer cachexia in the novel cancer cachexia mice model. Cancer cachexia muscle wasting symptom is initiated by abnormally high activin A, which activates the p38 MAPK pathway and further upregulates the transcription of E3 ligases, such as *Atrogin 1* (left panel). Skeletal muscle wasting progresses due to energy stress caused by hyper-metabolism of the growing tumor, which upregulates *FoxO3* transcription and further up-regulates E3 ligases such as *Atrogin 1* and *Murf1* through the AMPK pathway (right panel).

## Methods

### Mice

Transgenic mice with oocyte-specific expression of constitutively active PI3K (PIK3CA*) were generated as previously described(*22*). Homozygous female mice with a Cre-inducible knock-in allele for *Pik3ca\** were crossed with heterozygous *Gdf9-iCre* male mice (Jackson Laboratories, Bar Harbor, ME, USA). Cre+ female offspring overexpressed PI3KCA* and were referred to as Cre+ and were compared to Cre-littermate controls.

Female Cre-mice at 12 weeks of age were used for fasting experiments. One group of Cre-mice were fasted for 16 hours before euthanasia, while the other group of Cre-mice was fed normally. Male Cre-mice between 12 and 16 weeks of age were injected with 220 mg/kg of streptozotocin (572201, Millipore Sigma, St. Louis, MO, USA) to chemically induce irreversible type 1 diabetes. Non-fasting blood glucose levels were measured using the OneTouch Basic Glucose Monitor (Li LifeScan, AW, USA) and only those mice with a measurement of 450 mg/dL or greater were considered as type 1 diabetes mice. The mice were kept for 8-10 days after blood glucose increased until mice lost 20% of their body weight. PBS-injected Cre-male mice were used as the control for type 1 diabetes mice. Animals were housed and treated within Northwestern University’s Center of Comparative Medicine (Chicago, IL, USA) and were provided with food and water *ad libitum*. Temperature, humidity, and photoperiod (14 hrs light – 10 hrs dark) were kept constant.

### Measurement of body composition

The measurement of lean body mass and fat mass were conducted by quantitative nuclear magnetic resonance (qNMR; EchoMRITM, Houston, TX, USA). Whole-body MRI scan was conducted with the purpose of detecting tumor growth and assessing body composition and the distribution of fat tissue. All whole-body MRI measurements were conducted using a dedicated high-field (7Tesla) MRI (Bruker ClinScan). A 3-point Dixon MR sequence was then used to obtain images depicting only fat, only normal tissue, and both fat and normal tissue. All scans were conducted with mice under anesthesia using isoflurane mixed with oxygen (100% O_2_). Respiration and temperature were monitored using dedicated physiological monitoring devices (SA Instruments, NJ, USA).

### Histology and immunohistochemistry

The tibialis anterior muscle, liver, and stomach were collected from Cre- and Cre+ mice and fixed in Modified Davidson’s fixative (64133-10, Electron Microscopy Sciences, PA, USA) for 48 h at 4°C. Tissues were then processed and embedded in paraffin. Hematoxylin and eosin (H&E) staining was performed using standard methods. Serial sections (5 μm) were cut transversely through the TA muscle, liver, and stomach. TA sections were stained with anti-Laminin antibody (L9393, Millipore Sigma) for myofiber cross-sectional area (CSA) measurement(*5*). The quantification of CSA was performed using ImageJ (NIH software version 1.50i). Sections were also stained with anti-FOXO3 antibody (LS-B5284, LifeSpan Biosciences, Seattle, WA, USA) for cellular distribution of FOXO3.

### Blood glucose measurement

Cre+ mice with less than 5% weight loss (precachectic) or over than 15% weight loss but less than 20% weight loss (cachectic) and age-matched Cre-mice were used for non-fasting and fasting blood glucose measurement. Mouse tail veins were accessed with 31-gauge needle and a drop of blood was used for measuring blood glucose by the One Touch Basic Glucose Monitor system (Aviva).

### Glucose tolerance test

Cre+ mice with <5% weight loss (precachectic), Cre+ mice with 15% weight loss (cachectic), and age-matched Cre-mice were fasted for 16 h. Body weight and baseline blood glucose level were monitored for each mouse. Each mouse was fed 2 mg glucose/body weight using a 20% glucose saline solution. The mice were placed back into the cage and blood glucose levels were measured at each 15, 30, 60, and 90 min as described above.

### ^18^FDG PET/CT scan

Positron emission tomography-computed tomography (PET-CT) scanning was conducted using a ^18^F-fluorodeoxyglucose (FDG) tracer to monitor regional glucose uptake by tissues. All experiments were conducted on NanoScan8 PET-CT (Mediso, Budapest, Hungary). Cre+ mice and age-matched Cre-mice were deprived of food for 6 hours and *ad libitum* access to water before scanning. Fasting blood glucose levels were measured by OneTouch Basic Glucose Monitor system (Aviva) before glucose challenge. The mice were fed 20% glucose in saline solution at the dose of 2 mg/kg glucose, and then were intravenously injected with ^18^F-FDG tracer at a dose of ~10 μCi/g in 0.2 mL. Mice were kept awake and in a heated environment and their blood glucose levels were checked at 15, 30, and 60 min. At 60 min post-injection of ^18^F-FDG, mice were anesthetized in isoflurane mixed with 100% O_2_ and were placed in the holder inside PET/CT scanner. Body temperature and respiration were monitored continuously using the integrated physiological monitoring unit. Acquisition of a full body X-Ray CT scan (340 projections, 80 KeV, 340 msec exposure time) was initiated to generate a density map for attenuation correction during reconstruction of the PET images. The CT scan lasted 2 minutes and the final reconstructed spatial resolution was 250 μm in all three directions. Following the collection of the 3D tomographic data, the PET scan was initiated for a period of 20 minutes. After reconstruction, a comparison between different mice and different time points was done by converting the images to percent of injected dose (ID%). Representative figures of glucose uptake were analyzed by Mango (version 4.0.1, UTHSCSA) by choosing the sections at same locations from axial, coronal, and sagittal planes, respectively.

### ELISA

Blood was collected by cardiac puncture or tail blood draw. Activin A and GDF15 levels were determined by sandwich enzyme-linked immunosorbent assay (ELISA) kits for activin A (Anshlab, Webster, TX, USA) and GDF15 (R&D systems, Minneapolis, MN, USA). The limits of detection of these kits are 0.065 ng/mL and 2.2 pg/mL, respectively. IL-1β, IL-6, and TNF-α levels were measured using ELISA kits (ThermoFisher Scientific, Waltham, MA, USA), with sensitivities of <3 pg/mL, <7 pg/mL, and <3 pg/mL, respectively.

### Treatment

Recombinant Follistatin 288 (Fst288) was a gift from Dr. Thomas Thompson from The Ohio State University(*26*). Cre+ mice (n=6) were randomly assigned into two groups, one group of Cre+ mice were subcutaneously (s.c.) injected with 200 μg/kg Fst288 b.i.d, starting with mice that had 5% weight loss for 2 continuous days, which was considered to be the onset of cachexia-associated weight loss. Injections continued until the mice gained 5% body weight. The body weight was measured daily during the suspension of injection, and the second round of Fst288 injection was started when the Cre+ mice had lost approximately 10% of their body weight. The second round of Fst288 injection continued until Cre+ mice either gained over 5% body weight for at least 2 days or lost 15% of their body weight. Another group of Cre+ mice and a group of age-matched Cre-mice were s.c. injected with vehicle PBS based on the Fst288 injection schedule. Weight was recorded every day for each mouse. All vehicle-treated Cre+ mice were euthanized after 15% weight loss.

### Immunoblot analysis

The change in protein level from TA muscle was detected by immunoblot analysis. TA muscle tissues (10-20 mg per sample) were collected and homogenized in ice-cold lysis buffer (20 mM Tris-HCl, pH 8.0, 137 mM NaCl, 10% glycerol, 1% Triton-X, 2 mM EDTA) with protease inhibitor cocktail (04693116001, Roche Life Science, Indianapolis, IN, USA) and Phosphatase Inhibitor cocktail (P0044, Millipore Sigma, Burlington, MA,). Equal amounts (30 μg) of protein measured by the BCA protein assay kit (23225, ThermoFisher Scientific, Waltham, MA, USA) were loaded into a precast NuPAGE 4-12% gradient Bis-Tris gel. Electrophoresis was performed and proteins were dry transferred to a nitrocellulose membrane using an iBlot 2 system (ThermoFisher Scientific). The blots were probed with anti-phospho-AKT (Ser473) (9271) or anti-phospho-FOXO1 (Thr24) / FOXO3a (Thr32) antibody (9464) or anti-FOXO3a mAb (2497) or phospho-p38 MAPK (Thr180/Tyr182) (9211) or p38 MAPK (9212) or phospho-AMPKα (Thr172) (2535) or AMPKα (2532) (Cell Signaling Technology, Danvers, MA USA) or Insulin Receptor beta (ab69508) overnight at 4°C followed by anti-rabbit secondary antibody conjugated to horseradish peroxidase (ThermoFisher Scientific). Proteins were detected by Luminata Crescendo immunoblot HRP substrate (WBLUR0500Millipore Corporation, Burlington, MA, USA,) and exposed using a c500 imaging system (Azure Biosystems, Dublin, CA USA). The same blot was stripped with buffer (ThermoFisher Scientific) and re-probed with anti-α-tubulin antibody (Millipore Sigma), followed by anti-mouse secondary antibody conjugated to horseradish peroxidase (ThermoFisher Scientific). The proteins were exposed and detected using the c500 imaging system (Azure Biosystems). Protein quantification was performed by densitometry analysis on ImageJ (NIH software version 1.50i).

### Real-time quantitative PCR

Total RNA from TA muscle was extracted and purified using the RNeasy Fibrous Tissue Mini Kit (74704, QIAGEN, Germantown, MD). RNA samples were transcribed to cDNA by Superscript III First Strand Synthesis Supermix (18080-400, ThermoFisher Scientific). The real-time Taqman probes for Fbxo32 (*Atrogin1*, Mm00499523_m1), Trim63 (*Murf1*, Mm1185221_m1), Map1lc3a (*Lc3a*, Mm00458724_m1) and FoxO3 (*FoxO3*, Mm01185722_m1) were purchased from Taqman (ThermoFisher Scientific). The Rpl9 gene (*Rpl9*, Mm01208901_g1) served as an internal control gene for quantification. Real-time PCR was performed on a StepOnePlus real-time PCR system (Applied Biosystems, Life Technologies, CA, USA). The cycling conditions were a polymerase activation step (95°C for 10 min) and 40 cycles of PCR, including denaturation for 15 sec at 95°C and annealing-extension for 1 min at 60°C. Gene expression was analyzed using the comparative Ct (cycle threshold) method (ΔΔCt). The relative expression of the gene of interest was calculated by the following equation (using Atrogin1 in Cre+ muscle as an example): Atrogin1(Cre+) = 2(-ΔΔCt); ΔΔCt = [Ct (Atrogin1)-Ct (Rpl9)](Cre+) – [Ct (Atrogin1)-Ct (Rpl9)](Cre-).

### Statistical analysis

The results of ELISA, TA muscle CSA counting, RT-qPCR relative expression, immunoblot densitometry, and blood glucose are presented as mean ± SD. All of the graphs were generated by Prism 7 software (GraphPad Software Version 7, Inc., CA, USA). The data were analyzed by unpaired parametric t-tests with Welth’s correction (not assuming equal SDs) by Prism 7 software. The correlation data were analyzed using Spearman Correlation by Prism 7 software. P values of less than 0.05 were considered statistically significant.

## Acknowledgments

The authors thank Dr. Daniele Procissi in the Northwestern University Feinberg School of Medicine Preclinical Molecular Imaging Division @ Center for Translational Imaging for help with MRI and ^18^ FDG-PET/CT scanning, Dr. Weiming Song in the Northwestern University Comprehensive Metabolic Core for help with the EchoMRI experiment, and Dr. Stacey Tobin (The Tobin Touch, Inc.) for editorial assistance.

## Acknowledgements

Funding for this project was provided by the Watkins Chair of Obstetrics and Gynecology (T.K.W.), Cancer Center and the Lynn Sage Foundation through Northwestern Memorial Foundation, and the Center for Reproductive Health After Disease (P50 HD076188) from the National Centers for Translational Research in Reproduction and Infertility (NCTRI).

## Author contributions

Y.Z and J.Z. performed all the biological experiments, data analysis, and manuscript writing. S.Y.K. and T.K. provided the PI3K mice model. M.M.R and K.A.E maintained the mouse colony and performed genotyping. S.Y.K., T.K., and T.K.W. contributed to manuscript editing and critical review. T.K. provided intellectual contribution to the manuscript preparation. All authors approved the final version of the manuscript.

## References

1. M. J. Tisdale, Reversing cachexia. Cell 142, 511–512 (2010).

2. K. Fearon et al., Definition and classification of cancer cachexia: an international consensus. The Lancet. Oncology 12, 489–495 (2011).

3. K. Fearon, J. Arends, V. Baracos, Understanding the mechanisms and treatment options in cancer cachexia. Nature reviews. Clinical oncology 10, 90–99 (2013).

4. R. Ballarò, P. Costelli, F. Penna, Animal models for cancer cachexia. Current opinion in supportive and palliative care, (2016).

5. X. Zhou et al., Reversal of cancer cachexia and muscle wasting by ActRIIB antagonism leads to prolonged survival. Cell 142, 531–543 (2010).

6. S. Busquets et al., Myostatin blockage using actRIIB antagonism in mice bearing the Lewis lung carcinoma results in the improvement of muscle wasting and physical performance. Journal of cachexia, sarcopenia and muscle 3, 37–43 (2012).

7. M. D. Gomes, S. H. Lecker, R. T. Jagoe, A. Navon, A. L. Goldberg, Atrogin-1, a muscle-specific F-box protein highly expressed during muscle atrophy. Proc Natl Acad Sci U S A 98, 14440–14445 (2001).

8. O. Rom, A. Z. Reznick, The role of E3 ubiquitin-ligases MuRF-1 and MAFbx in loss of skeletal muscle mass. Free radical biology & medicine, (2015).

9. A. M. J. Sanchez, R. B. Candau, H. Bernardi, FoxO transcription factors: their roles in the maintenance of skeletal muscle homeostasis. Cellular and molecular life sciences: CMLS 71, 1657–1671 (2014).

10. M. Sandri et al., Foxo transcription factors induce the atrophy-related ubiquitin ligase atrogin-1 and cause skeletal muscle atrophy. Cell 117, 399–412 (2004).

11. C. Mammucari et al., FoxO3 controls autophagy in skeletal muscle in vivo. Cell metabolism 6, 458–471 (2007).

12. L. M. Bollinger, C. A. Witczak, J. A. Houmard, J. J. Brault, SMAD3 augments FoxO3-induced MuRF-1 promoter activity in a DNA-binding-dependent manner. American journal of physiology. Cell physiology 307, C278–287 (2014).

13. J. A. Carson, K. A. Baltgalvis, Interleukin 6 as a Key Regulator of Muscle Mass during Cachexia. Exercise and Sport Sciences Reviews 38, 168–176 (2010).

14. H. J. Patel, B. M. Patel, TNF-alpha and cancer cachexia: Molecular insights and clinical implications. Life Sci 170, 56–63 (2017).

15. M. M. Matzuk et al., Development of cancer cachexia-like syndrome and adrenal tumors in inhibin-deficient mice. Proceedings of the National Academy of Sciences of the United States of America 91, 8817–8821 (1994).

16. J. L. Chen et al., Elevated expression of activins promotes muscle wasting and cachexia. The FASEB journal: official publication of the Federation of American Societies for Experimental Biology, (2014).

17. T. B. Dschietzig, Myostatin - From the Mighty Mouse to cardiovascular disease and cachexia. Clin Chim Acta 433, 216–224 (2014).

18. L. Lerner et al., MAP3K11/GDF15 axis is a critical driver of cancer cachexia. Journal of cachexia, sarcopenia and muscle 7, 467–482 (2016).

19. E. Lach-Trifilieff et al., An antibody blocking activin type II receptors induces strong skeletal muscle hypertrophy and protects from atrophy. Molecular and cellular biology 34, 606–618 (2014).

20. J. Zhu et al., Virtual High-Throughput Screening To Identify Novel Activin Antagonists. Journal of medicinal chemistry, (2015).

21. M. I. Polkey et al., Activin Type II Receptor Blockade for Treatment of Muscle Depletion in Chronic Obstructive Pulmonary Disease. A Randomized Trial. Am J Respir Crit Care Med 199, 313–320 (2019).

22. S.-Y. Kim et al., Cell autonomous phosphoinositide 3-kinase activation in oocytes disrupts normal ovarian function through promoting survival and overgrowth of ovarian follicles. Endocrinology 156, 1464–1476 (2015).

23. S.-Y. Kim et al., Constitutive Activation of PI3K in Oocyte Induces Ovarian Granulosa Cell Tumors. Cancer research 76, 3851–3861 (2016).

24. W. Chen, T. K. Woodruff, K. E. Mayo, Activin A-induced HepG2 liver cell apoptosis: involvement of activin receptors and smad proteins. Endocrinology 141, 1263–1272 (2000).

25. K. C. H. Fearon, D. J. Glass, D. C. Guttridge, Cancer cachexia: mediators, signaling, and metabolic pathways. Cell metabolism 16, 153–166 (2012).

26. T. B. Thompson, T. F. Lerch, R. W. Cook, T. K. Woodruff, T. S. Jardetzky, The structure of the follistatin:activin complex reveals antagonism of both type I and type II receptor binding. Developmental cell 9, 535–543 (2005).

27. H. Ding et al., Activin A induces skeletal muscle catabolism via p38beta mitogen-activated protein kinase. J Cachexia Sarcopenia Muscle 8, 202–212 (2017).

28. K. Nakashima, Y. Yakabe, AMPK activation stimulates myofibrillar protein degradation and expression of atrophy-related ubiquitin ligases by increasing FOXO transcription factors in C2C12 myotubes. Bioscience Biotechnology and Biochemistry 71, 1650–1656 (2007).

29. D. G. Hardie, AMPK: positive and negative regulation, and its role in whole-body energy homeostasis. Current Opinion in Cell Biology 33, 1–7 (2015).

30. G. L. Rohdenburg, A. Bernhard, O. Krehbiel, Sugar tolerance in cancer. Journal of the American Medical Association 72 1528–1530 (1919).

31. A. Amrani et al., Glucose homeostasis in the nonobese diabetic mouse at the prediabetic stage. Endocrinology 139, 1115–1124 (1998).

32. L. Lerner et al., Plasma growth differentiation factor 15 is associated with weight loss and mortality in cancer patients. Journal of cachexia, sarcopenia and muscle 6, 317–324 (2015).

33. P. J. Emmerson et al., The metabolic effects of GDF15 are mediated by the orphan receptor GFRAL. Nature medicine 23, 1215–1219 (2017).

34. S. M. Judge et al., Genome-wide identification of FoxO-dependent gene networks in skeletal muscle during C26 cancer cachexia. BMC cancer 14, 997 (2014).

35. N. Johns et al., Clinical classification of cancer cachexia: phenotypic correlates in human skeletal muscle. PloS one 9, e83618 (2014).

36. L. De Angelis, S. Balasubramanian, L. Berghella, Akt-mediated phosphorylation controls the activity of the Y-box protein MSY3 in skeletal muscle. Skeletal muscle 5, 18 (2015).

37. G. Pallafacchina, E. Calabria, A. L. Serrano, J. M. Kalhovde, S. Schiaffino, A protein kinase B-dependent and rapamycin-sensitive pathway controls skeletal muscle growth but not fiber type specification. Proceedings of the National Academy of Sciences of the x x xUnited States of America 99, 9213–9218 (2002).

38. T. K. Woodruff, Role of inhibins and activins in ovarian cancer. Cancer treatment and research 107, 293–302 (2002).

39. B. T. O’Neill et al., FoxO Transcription Factors Are Critical Regulators of Diabetes-Related Muscle Atrophy. Diabetes 68, 556–570 (2019).

40. I. Bosaeus, P. Daneryd, E. Svanberg, K. Lundholm, Dietary intake and resting energy expenditure in relation to weight loss in unselected cancer patients. Int J Cancer 93, 380–383 (2001).

