## Supplemental Figures and Legends for "Stage-specific muscle wasting mechanisms in a novel cancer cachexia mouse model for ovarian granulosa cell tumor"

**Supplemental Figure 1.** (**A**) Breeding strategy for producing mice with constitutive PI3K activation in early-stage oocytes. (**B**) H&E staining of representative ovaries from a Cre- mouse (above) and a Cre+ mouse (below) on PD60. The ovarian follicles were only in Cre- mice (blue arrows) and not in Cre+ mice. Scale bar = 100μm. (**C**) MRI image of ovarian tumors (red arrows) in Cre+ mouse (right) on PD60 and age-matched Cre- mouse (left). (**D**) Size of tumor ($mm^3$) quantitated by MRI after PD50 (day 0). (**E**) Survival curve of Cre+ and Cre- mice after PD60 (n=13). (**F**) Representative TA muscle on the hind limbs of Cre+ mice after 15% weight loss (right) and age-matched Cre- mice (left). Tibialis anterior (TA) muscle are indicated by the black arrows. (**G**) Images of abdomen from Cre+ mouse and age-matched Cre- mouse. Tumor is shown in Cre+ mice (red arrow) and the fat pad disappears in the Cre+ mouse compared with age-matched Cre- mouse (blue arrow). (**H**) MRI images of abdomen fat pad in a Cre- mouse (blue arrow) and Cre+ mouse (red arrow).

**Supplemental Figure 2.** (**A**) RT-PCR analysis of *Caspase 3* in TA muscle collected from Cre+ mice with 15% weight loss and age-matched Cre- mice (n=3). (**B**) Serum TNF-α levels in Cre+ mice with weight loss from 5% to 20% (n=8). Non-paired t-test was used for statistical analysis by Prism 7 software. Data are shown as mean ± s.d. of biological replicates. * and ** represent $P<0.05$ and $P<0.01$, respectively.

**Supplemental Figure 3.** (**A**) Percentage of body weight recorded daily for Cre+ mice with PBS injection (n=3). (**B**) Percentage of body weight recorded daily for Cre+ mice with Fst288 injection (n=3). (**C**) Correlation of *Murf1* mRNA expression and weight loss in Cre+ mice (n=16). (**D**) Correlation of *Atrogin1* mRNA expression and weight loss in Cre+ mice (n=16). (**E**) Correlation of *Lc3* mRNA expression and weight loss in Cre+ mice (n=8). Statistical analysis was done by Spearman correlation using Prism 7 software. *, and ** represent $P<0.05$ and $P<0.01$, respectively.

**Supplemental Figure 4.** (**A**) RT-PCR analysis of *FoxO3* in skeletal TA muscle collected from Cre- mice with 16h fasting (n=3) and no fasting control (n=3). (**B**) RT-PCR analysis of *FoxO3* in skeletal TA muscle collected from Cre- mice with STZ-induced diabetes (n=3) and vehicle injection control (n=3). (**C**) Representative image of immunoblot of p-FOXO3, FOXO3, and α-tubulin protein for the TA muscle lysate collected from Cre- mice with fasting and non-fasting controls, as well as Cre- mice with STZ-induced diabetes and vehicle injection control. (**D-E**) Quantification of densitometry analysis for immunoblot images for protein ratio of (**D**) p-FOXO3/FOXO3 and (**E**) FOXO3/α-tubulin in Cre- mice with fasting and non-fasting control (n=3). (**F-G**) Quantification of densitometry analysis for immunoblot images for protein ratio of (**F**) p-FOXO3/FOXO3 and (**G**) FOXO3/α-tubulin in Cre- mice with STZ-induced diabetes (n=3) and vehicle injection control (n=3) using ImageJ software. Non-paired t-test was used for statistical analysis by Prism 7 software. Data are shown as mean ± s.d. of biological replicates. *, **, and *** represent $P$<0.05, $P$<0.01, and $P$<0.005, respectively.

**Supplemental Figure 5.** (**A**) Representative immunoblot of insulin receptor β, p-AKT, AKT, and α-tubulin protein in TA muscle lysates collected from Cre+ mice with 15% weight loss and age-matched Cre- mice. (**B**) Quantification of glucose uptake by brown fat tissue in Cre+ mice (n=3) at the precachexia stage and (n=3) age-matched Cre- mice using $^{18}$F-FDG as tracer with the MRI scan images. Unpaired *t*-test was used to test statistical significance by Prism 7.

**Supplemental Figure 6.** (**A**) Representative immunoblots of p-p38 MAPK, p-38 MAPK and α-tubulin protein in TA muscle lysates collected from Cre+ mice with 5% weight loss, Cre+ mice with 15% weight loss, and age-matched Cre- mice (n=3). (**B**) Representative immunoblots of p-FOXO3, FOXO3, and α-tubulin protein in TA muscle lysates collected from Cre+ mice with 5% weight loss, Cre+ mice with 15% weight loss, and age-matched Cre- mice (n=3). (**C**)

Representative immunoblots of AMPKα, p-AMPKα, and α-tubulin protein in TA muscle lysates collected from Cre+ mice with 5% weight loss, Cre+ mice with 15% weight loss, and age-matched Cre- mice (n=3).

### Supplemental Figure 1

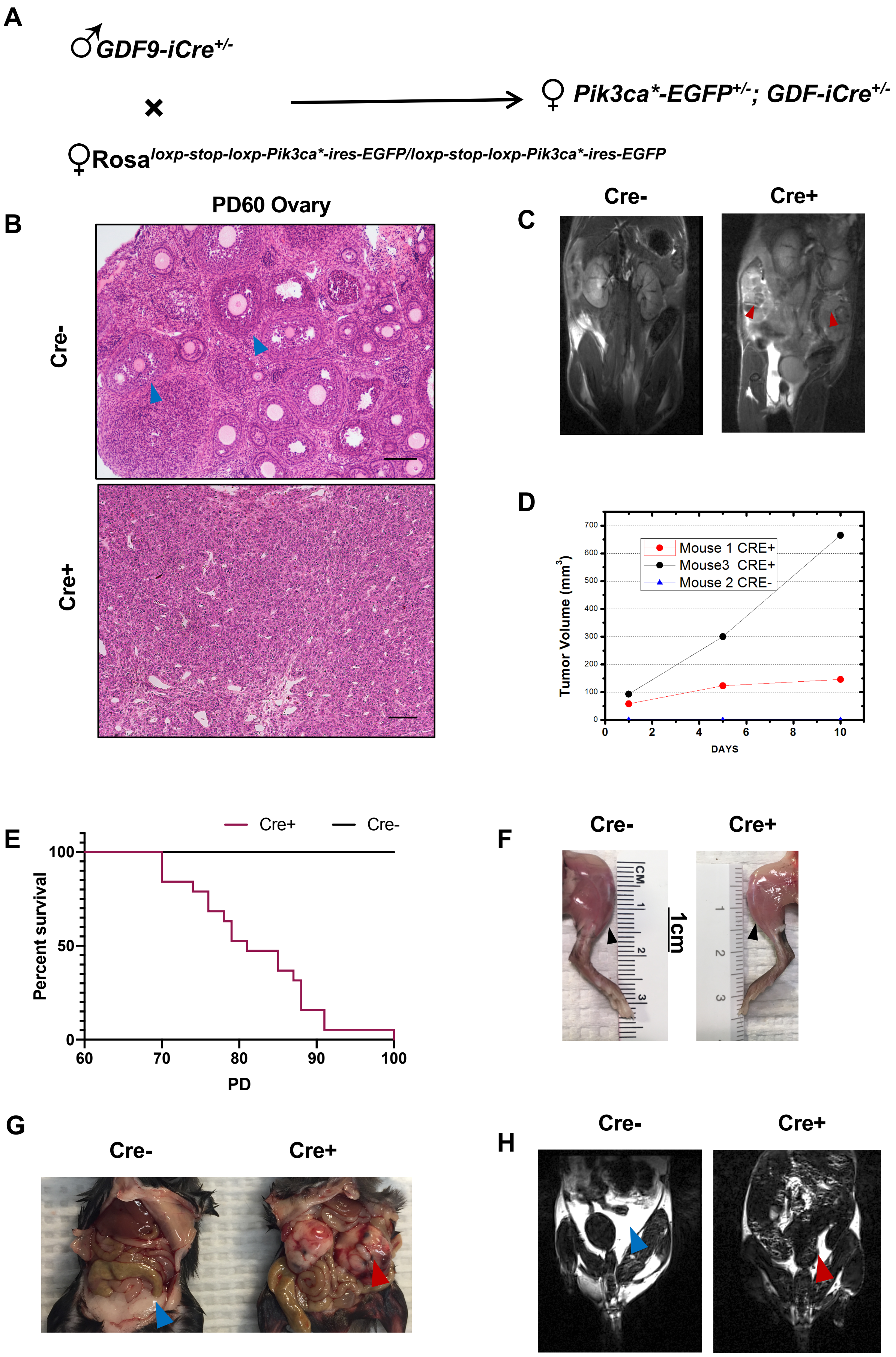

**Supplemental Figure 2**

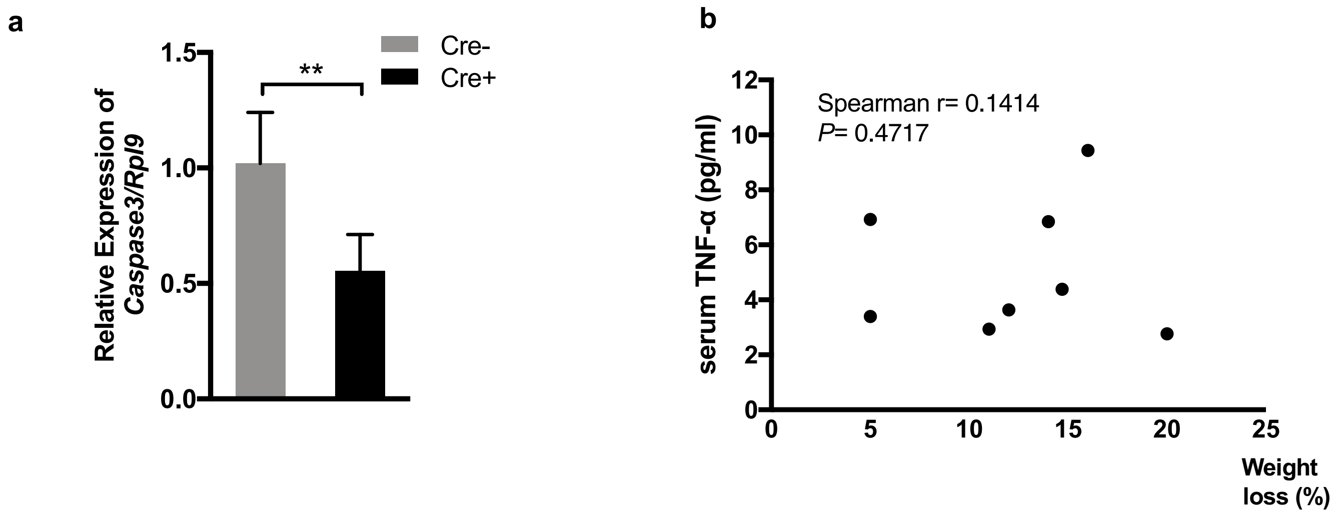

### Supplemental Figure 3

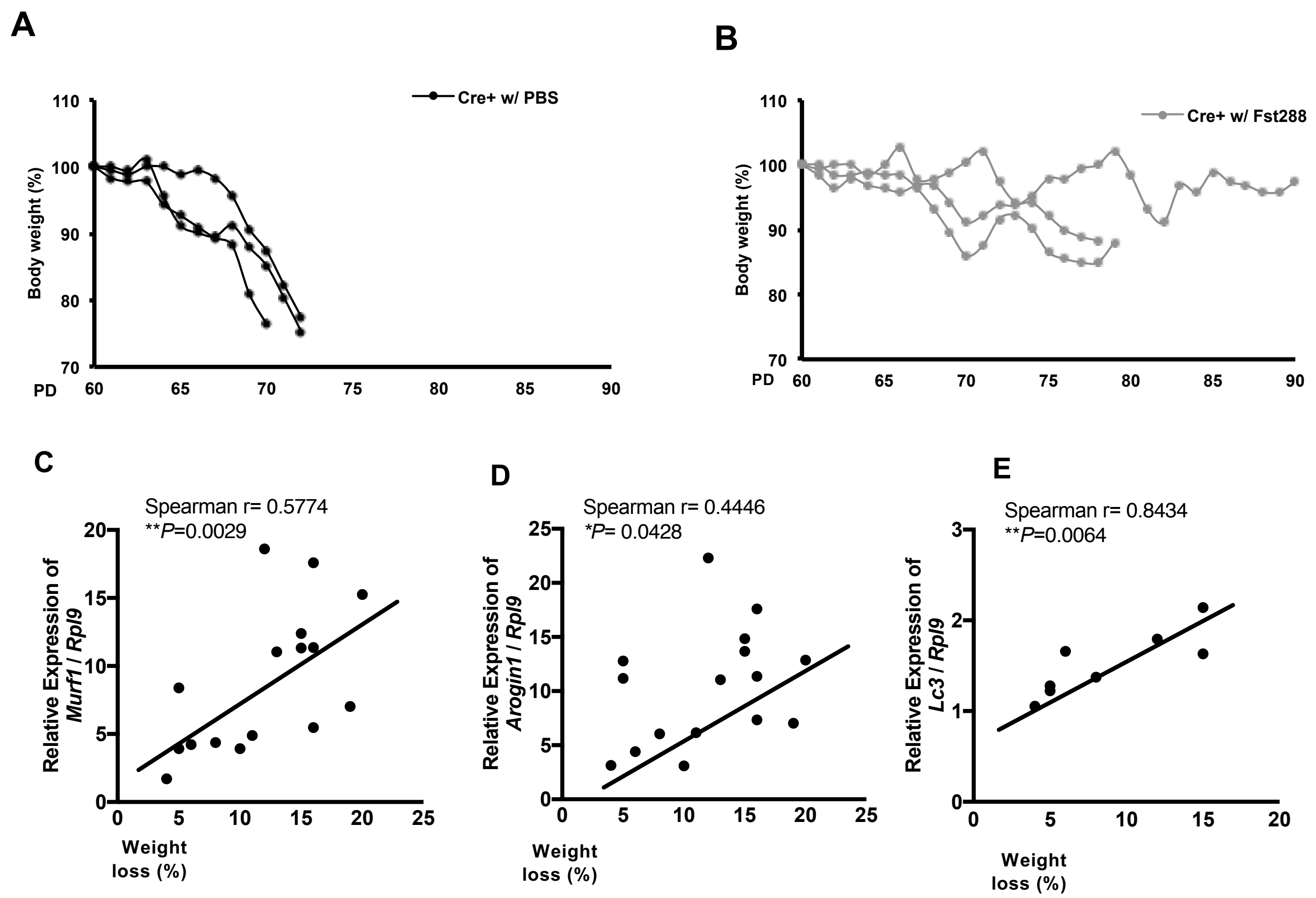

### Supplemental Figure 4

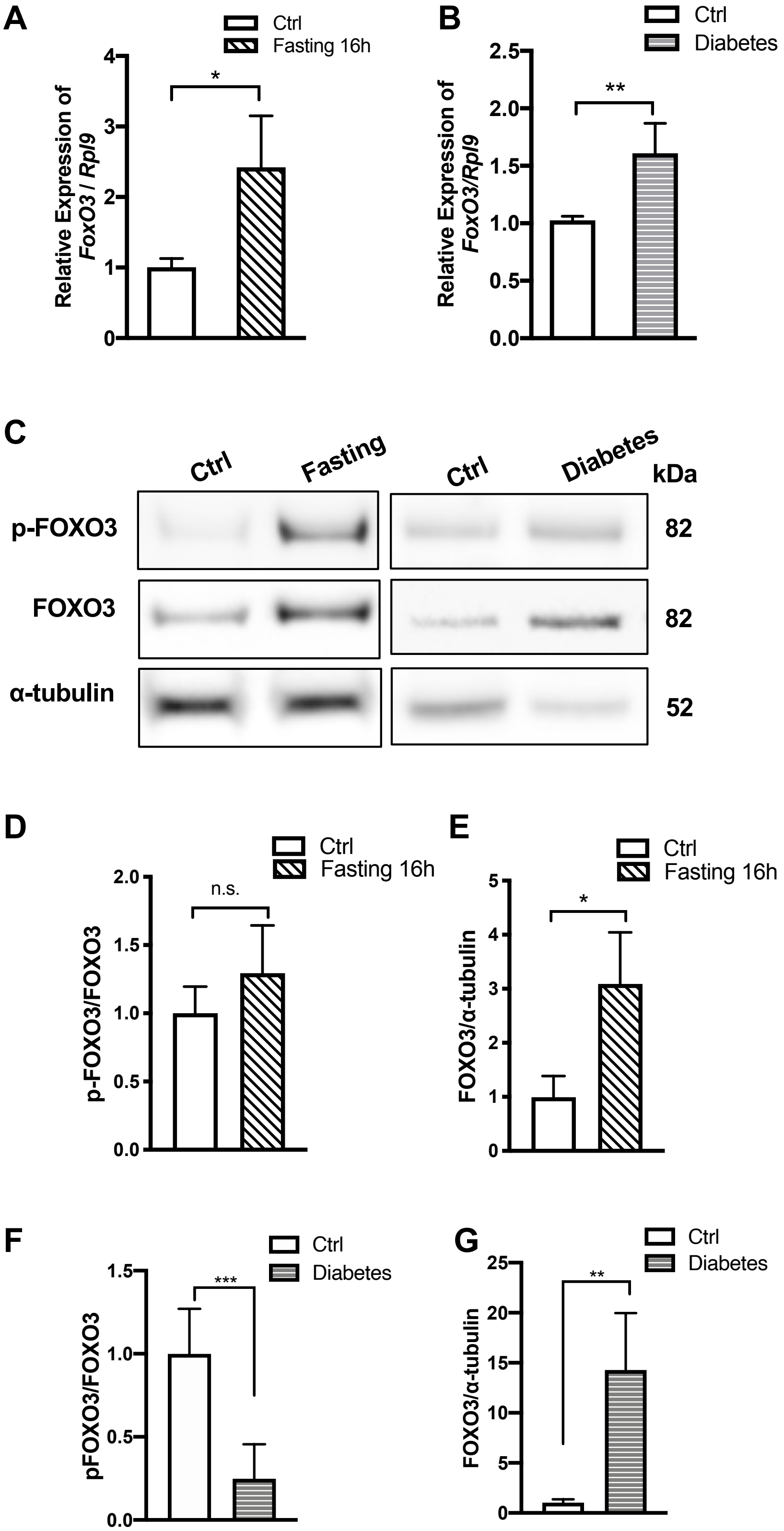

### Supplemental Figure 5

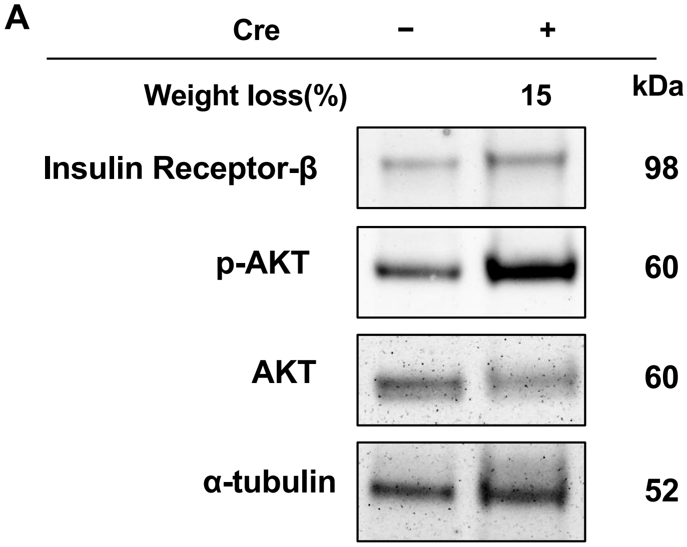


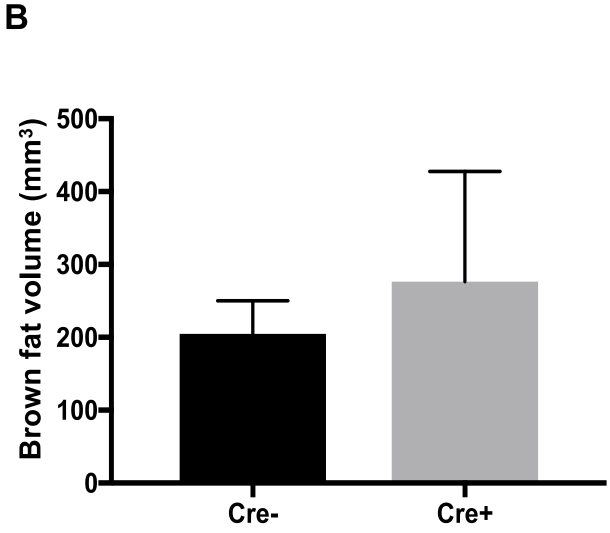

### Supplemental Figure 6

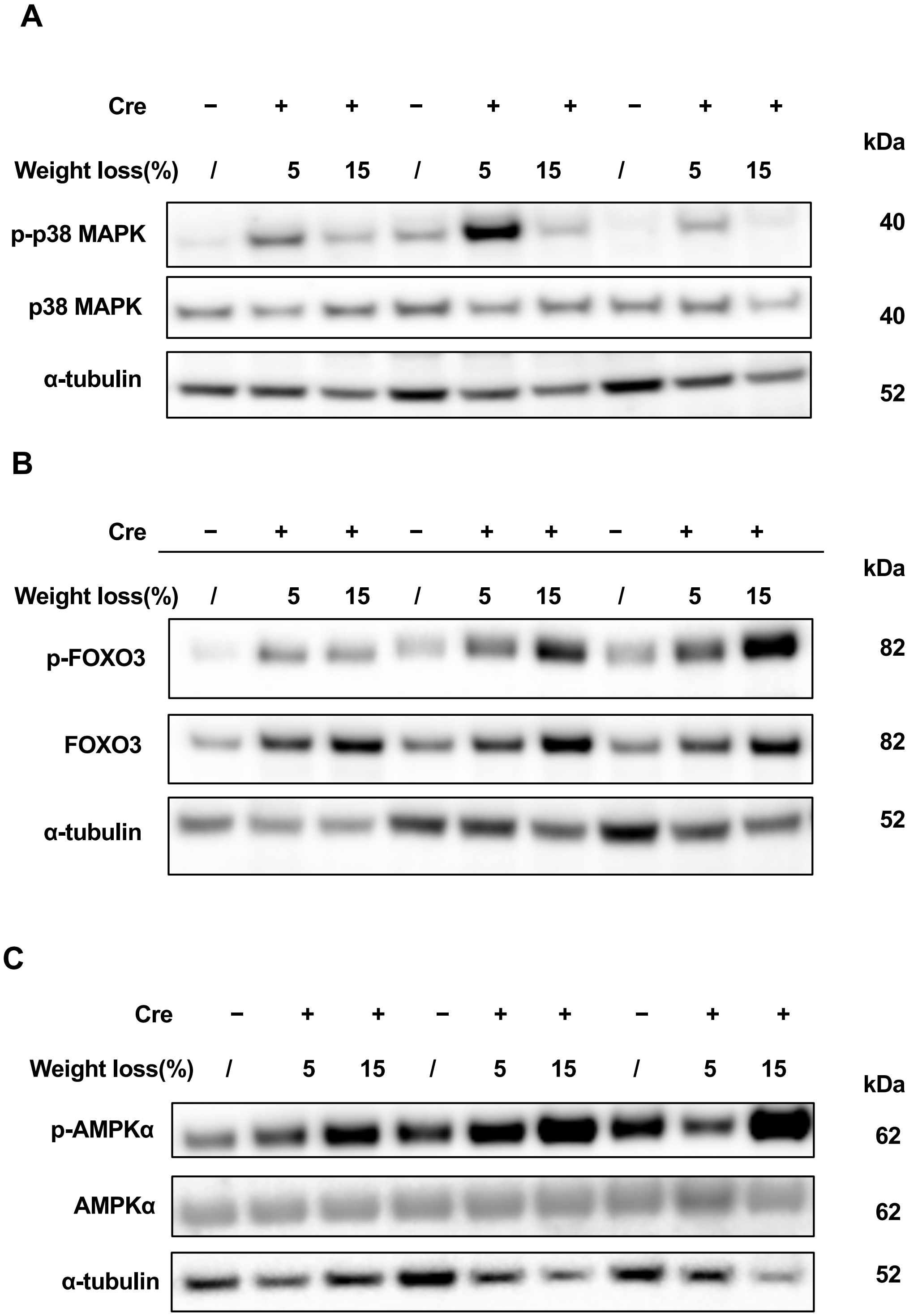
